# Co-evolution of human influenza A and Epstein Barr virus-specific CD8 ex vivo memory T cell receptor BV repertoires with increasing age

**DOI:** 10.1101/2022.12.15.520600

**Authors:** Fransenio Clark, Anna Gil, Nuray Aslan, Dario Ghersi, Liisa K. Selin

**Affiliations:** Department of Pathology, University of Massachusetts Medical School, Worcester, MA, USA; University of Massachusetts Medical School, Worcester, MA, USA; School of Interdisciplinary Informatics, University of Nebraska at Omaha, NE, USA

**Keywords:** Influenza A virus, IAV, Epstein Barr virus, EBV, repertoire, T cell receptor, TCR

## Abstract

CD8 memory T cells are generated during primary infection with intracellular pathogens, such as viruses. These cells play an important role in the protection of the host upon re-infection with the same pathogen. In this study, we compare CD8 memory T cell receptor (TCR) BV repertoires directly ex vivo for two common human viruses, influenza A virus (IAV), an RNA virus that frequently re-infects due to a high rate of genetic mutation, and Epstein-Barr virus (EBV), a DNA virus, which persists in B cells for life, in the 95% of people that become infected. In cross-sectional and longitudinal studies of EBV seropositive, HLA-A2+, young (18-22 years), middle age (25-59 years), and older (>60 years) donors, we demonstrate that CD8 memory TCR repertoires to three immunodominant epitopes, known to have cross-reactive responses, IAV- M1_58-66_ (M1), EBV-BMLF1_280-288_ (BM), and EBV-BRLF_109-117_ (BR) co-evolve as individuals age. Cross-sectional studies showed that IAV-M1- and both EBV-specific repertoires narrowed their TRBV usage with increasing age manifesting to different degrees for each epitope. In fact, narrowing of EBV-BM and EBV-BR-specific TRBV usage correlated with increasing age. IAV- M1-specific TRBV usage was significantly narrowed by middle-age. There was evidence that TRBV usage was changing with increasing age. For instance, IAV-M1-specific dominant BV19 usage appeared to become bimodal showing either high or low frequency of usage in the older age group, while BV30 usage frequency directly correlated with age. For the EBV epitope-specific responses there was preferential usage of particular TRBV and changes in the hierarchy of BV family usage in the different age groups. There appeared to be focusing of the TRBV repertoire by all 3 epitopes to three common BV in the older donors, which would be consistent with retention of crossreactive TCR suggesting co-evolution. Longitudinal studies tracking two donors over 14- 15 years (middle age to older) showed that there were continuous modulations in the TCR repertoire of IAV-M1, EBV-BM and EBV-BR-specific responses over time. There was evidence that acute IAV infection could contribute to these changes in TCR repertoire. This could be occurring by the TCR cross-reactivity that is known to exist between these 3 epitopes, and which appeared to be enhanced during acute IAV infection based on increased usage of common shared TRBV. These studies suggest that virus-specific TCR repertoires change over time as individuals’ age leading to narrowing of the repertoire favoring retention of potentially crossreactive TCR.

## Introduction

CD8 T-cell recognition of virus-infected cells requires a specific interaction between short peptides presented by HLA Class I molecules on infected cell surfaces and TCRαβ heterodimers on CD8 T-cells. CD8 T-cell TCR repertoires to common viruses (IAV, EBV, CMV) are highly diverse and individualized i.e. “private”. However, despite this diversity there are clonotypes with “public” features, (*i.e.* preferential usage of particular V region or conserved or identical amino acid motifs within the CDR3α/β (1) for each epitope that appear to be favored for expansion, likely due to selection for optimal structural interactions (2, 3). Disease etiology and diagnosis by TCR repertoire analysis is beginning to gain more attention as technology improves (37).

Influenza is a single-stranded RNA virus, that contains 3 subtypes, Type A (IAV), B (IBV), and C (ICV). IAV is the most important subtype, as it is capable of causing both an epidemic and pandemic, such as, the 1918 Spanish influenza A pandemic (4, 5). Accumulating mutations and re-assortments of certain viral gene segments result in generating new modified virus strains. The new modified viruses can be closely related to their ancestors due to minor genetic mutations known as antigenic drift, or distantly related as the result of major genetic changes following the recombination/reassortment of 2 different viruses known as antigenic shift (6, 7). Most immunocompetent individuals, that become infected with IAV will experience an acute illness, defined by fever, body aches, fatigue, and coughs, which resolves within two weeks (8). Each year, influenza vaccines targeting seasonal strains are developed to prevent infection and deadly secondary complications, such as bacterial pneumonia (9, 10). Despite the availability of a vaccine, influenza is a highly infectious and lethal virus that has infected 35.5 million, led to the hospitalization of more than 490,000, and killed 34,200 individuals around the world during the 2018-2019 season (11). Older individuals, those >50 years old, are particularly at risk and accounted for greater than 90% (>31,000) of the deaths during the 2018-2019 season (11–13).

Epstein Barr virus (EBV) is a double-stranded DNA virus that infects up to 95% of the world’s population by the 4^th^ decade of life. EBV infects oropharyngeal epithelium cells, then takes up latency in resting B cells for life (14–18). It is responsible for significant immune-pathology including acute infectious mononucleosis (AIM), a disease caused by a massive CD8 T cell proliferation leading to severe prolonged fatigue, lymphadenopathy, sore throat, and splenomegaly (19). EBV infection is associated with several other diseases such as Hodgkin’s lymphoma, Burkitt lymphoma, nasopharyngeal carcinoma as well as autoimmune disorders, such as multiple sclerosis, systemic lupus erythematosus, rheumatoid arthritis, and Sjögren’s syndrome (20, 21). Older donors have been observed to have a significantly higher viral load in peripheral blood B cells, plasma, early antigen and viral capsid antigen titers compared to middle-age donors, suggesting that older donors are having difficulty controlling this persistent virus (22) perhaps putting them at greater risk for all the diseases associated with this virus.

CD8 T cells are extremely important in providing resistance to new strains of IAV during infection and in keeping persistent EBV infection under control (23–29). For instance, several lines of evidence suggest that EBV-specific CD8 T-cells are important for the control of EBV long term (30), including successful treatment of EBV-associated lymphoproliferative disorders, post-transplant associated EBV infections and multiple sclerosis which has been associated with poor EBV control (31), by adoptive transfer of EBV-specific CD8 T-cells (32, 33). However, as individuals age, the immune system becomes increasingly dysfunctional, a phenomenon known as immunosenescence. Immunosenescence is characterized by dramatic changes in cell intrinsic and extrinsic factors including reduced lymphocyte development, proliferation, cytotoxicity, and repertoire diversity caused by thymic involution. These changes rapidly increase with age and are significant in impairing the ability of CD8 T cells to fight viral infections, leading older adults to becoming more susceptible to infections (34–41). A recent paper by Mark Davis and colleagues, highlights the importance of understanding how the immune system ages. They use an immune age scoring system based on using immune cellular phenotyping, which may provide a more accurate measure of susceptibility to subsequent infections than just chronological age (42).

Studies have shown that following the rapid increase in thymic involution after birth, rates steady to 3% involution each year until middle-age, and then 1% with each following year (43). As thymic involution greatly impacts the TCR repertoire diversity, many studies have focused on changes to TCR diversity in the total naive and memory phenotype of CD8 T cell populations in aging humans and mice (young vs. older) (44–50). However, there are few if any reports where changes of human antigen-specific TCR repertoire in 3 age groups, from young to older, simultaneously, was examined. Also, antigen-specific TCR repertoires of healthy donors as young as 18-19 years old are understudied. We were particularly interested in tracking the changes of the TCR repertoire to IAV and EBV-specific epitopes that are known to be highly conserved, protective, well-defined, and immunodominant in 100% of the HLA-A2.01+ donors (51). These epitopes include IAV- M1_58-66_ (M1), a matrix protein epitope from IAV, and EBV-BMLF1_280-288_ (BM) and EBV- BRLF1_109-117_ (BR), early and immediate early lytic protein epitopes, respectively, of EBV. EBV- BM and EBV-BR epitope-specific responses account for up to 20% of EBV-specific responses during acute and persistent infection (19, 52–55). Public TRBV features have been well defined for these three epitopes largely in middle-aged donors or patients with AIM. For instance, IAV- M1 TCR repertoires have highly increased usage of BV19 (56–59), while EBV-BM TCR repertoires use a number of different public TRBV usually with 2 of them being dominant in one donor that vary from person to person including BV20, BV14, BV2, BV29 and BV9 (60–63). The EBV-BR TCR repertoire appears to be selected in AIM donors by a dominant usage of TRAV8.1, associated with multiple different TRBV often with 5-6 different BV in anyone donor with a great deal of variability between donors i.e. private specificity (60, 61, 64). Responses to these three immunodominant epitopes are also known to be highly cross-reactive particularly during AIM (65–67). Here, we will track the IAV-M1, EBV-BM and EBV-BR epitope specific TRBV repertoires to better understand how they change with increasing age and potentially co-evolve due to TCR crossreactivity.

## Results

### Study populations

For these studies, we recruited and enrolled 40 healthy, IAV-and EBV-immune, HLA-A2.01+ donors. We used 3 age groups defined as young (18-22 years old) (YSP), middle age (25-59 years old) (MSP), and older (>60 years old) donors seropositive to EBV (OSP) (Supplemental Table.1). The average age of young donors was 19 years old, 37 years old for middle-age and 74 years old for older donors. These age ranges were chosen based on general convention and our previous published (56, 57) observations on changes in TCR repertoire with age.

*IAV-M1, EBV-BM and EBV-BR frequencies in different age groups.* Prior to studying TRBV repertoire we wanted to establish if there were any major differences in the frequency of IAV-M1, EBV-BM and EBV-BR-specific cells in these three different age groups by ex vivo tetramer/dextramer staining in PBMC. *Ex vivo* IAV-M1 and EBV-BR tetramer positive CD8 T cell frequencies were similar in all three age groups, indicating a relatively stable population with increasing age (Figure S1). In contrast, there was a 4-fold increase in EBV-BM-tetramer positive CD8 T cell frequency from young to middle age with a subsequent a 2-fold decrease in older donors suggesting that this response may be susceptible to change with increasing age. This change in EBV-BM frequency with age is consistent with previous reports of poorly understood fluctuations in frequency of EBV-BM-specific CD8 T cell responses with age (22) and CMV seropositivity (68).

### Evidence that epitope-specific TRBV diversity changes with increasing age

When we directly compared the diversity of the TRBV usage of each epitope-specific response between the three different age groups using Simpson Diversity Index (SDI), the middle age donors had significantly decreased IAV-M1 TRBV diversity compared to young adults (Figure 1A). There was a similar trend in narrowing of the TRBV repertoire diversity in the EBV-BM and EBV-BR responses by middle age, but most likely due to sample size this did not quite reach significance (Figure 1A). We therefore questioned whether there was any evidence of an inverse correlation between the frequency of different TRBV usage and age consistent with TRBV repertoire usage narrowing with increasing age. We found that both EBV-BM and EBV-BR TCR repertoire diversity as measured by SDI inversely correlated with increasing age (Figure 1Bii,iii). This was not, however, true for the IAV-M1 response, which appeared to have more complex changes occurring in the older age group where many donors had developed broad TRBV repertoires (Figure 1Bi).

**Figure 1.**
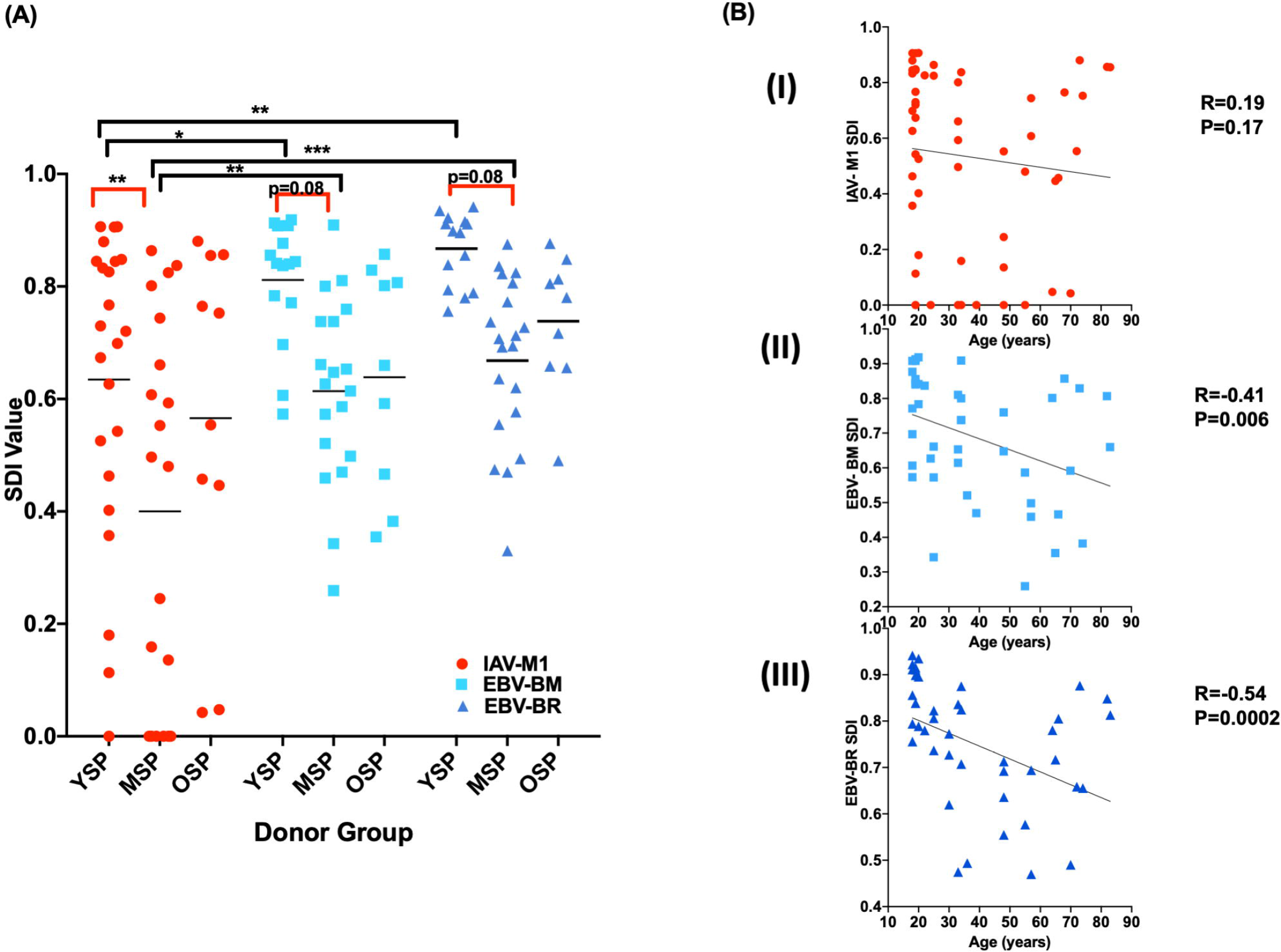
IAV-M1, EBV-BM, and EBV-BR TRBV repertoire diversity differed between age groups and between the epitope-specific responses within each age group. Ex vivo sorted CD8 T cells freshly isolated from peripheral blood of donors were co-stained with IAV-M1, EBV-BM and EBV-BR specific tetramers and TCR Vβ monoclonal antibodies. The Simpson Diversity Index (SDI) value was used to assess TRBV diversity and calculated by using the frequency of different TRBV measured in *ex vivo* IAV-M1, EBV-BM and EBV-BR tetramer+ CD8 T cells in each individual donor (YSP, n=14-23; MSP, n=19-20, OSP, n=9-10). A) Middle-age donors have decreased IAV-M1, EBV-BM and EBV-BR specific TRBV diversity compared to young donors. In young donors and middle-aged donors, the IAV-M1-specific TRBV usage was significantly less diverse than in EBV-BM and EBV-BR-specific responses. Multi-variant ANOVA with adjusted p values for multiple comparisons: * p<0.05, **p<0.01, ***p<0.001, ****p<0.0001. B). EBV-BM and EBV-BR TRBV repertoire diversity decreased with increasing age. A Spearman correlation between (i) IAV-M1, (ii) EBV-BM and (iii) EBV-BR specific TCR Vβ repertoire diversity as assessed by SDI and increasing age (years) was performed using all donors. IAV-M1 SDI shown in red, EBV-BM SDI shown in turquoise, and EBV-BR SDI shown in dark blue.

The pattern of TRBV repertoire diversity between the three different epitope-specific responses within each age group also changed (Figure 1A). IAV-M1-specific TRBV repertoire diversity was decreased compared to EBV-BM or -EBV-BR in both young and middle age groups, but in the older group this difference had disappeared, IAV-M1 TRBV diversity was no longer significantly less than EBV-BM or EBV-BR (Figure 1A). This may be explained by the fact that while EBV-BM and EBV-BR TRBV repertoires are getting narrower in older donors by direct correlation studies (Fig 1A,Bii, iii), the mean TCR repertoire diversity of IAV-M1-specific responses appeared to be increasing in some older donors as compared to middle-aged donors (Figure 1B).

Altogether, these data suggest that IAV-M1, EBV-BM and EBV-BR specific TRBV repertoire diversity was narrowing with increasing age, but this trend manifests to different degrees with each epitope response. The difference in diversity between epitope-specific responses, i.e. IAV-M1 is narrower than EBV-BM or EBV-BR, disappears in the older age group as it appears IAV-M1 repertoire is getting broader at least in some individuals, while the EBV-epitope specific responses continue to narrow.

### Antigen-specific TRBV monoclonal antibody staining: a highly reproducible technique

Co-staining with epitope-specific tetramers and TRBV monoclonal antibodies (mAbs) is a useful screening technique (as compared to TCR deep sequencing) to examine changes in TCR repertoire when examining a large population, particularly when examining 3 different epitope-specific responses in each individual. To assess the reproducibility of this method, we co-stained freshly isolated CD8 T cells in triplicate from the same individual with IAV-M1 and EBV-BM and -BR tetramers and TRBV repertoire mAbs. In a representative donor, the triplicate set of IAV- M1 tetramer co-staining with mAbs revealed similar frequencies in TRBV19, TRBV6.5-6.7, and TRBV27 family usage (Pearson’s r=0.93-1.00 and p<0.0001); triplicate staining of EBV-BM tetramer+ cells demonstrated almost identical frequencies of TRBV14 and TRBV27 (Pearson’s r=1.00 and p<0.0001); triplicate staining of EBV-BR tetramer+ cells showed similar frequencies of TRBV19 and TRBV6.5-6.7 usage (Pearson’s r=0.90-1.00, p<0.0001) (Figure 2). These data suggest that the BV mAb staining of tetramer positive cells is an accurate and reproducible technique that can show differences in repertoire usage in the different epitope-specific responses.

**Figure 2.**
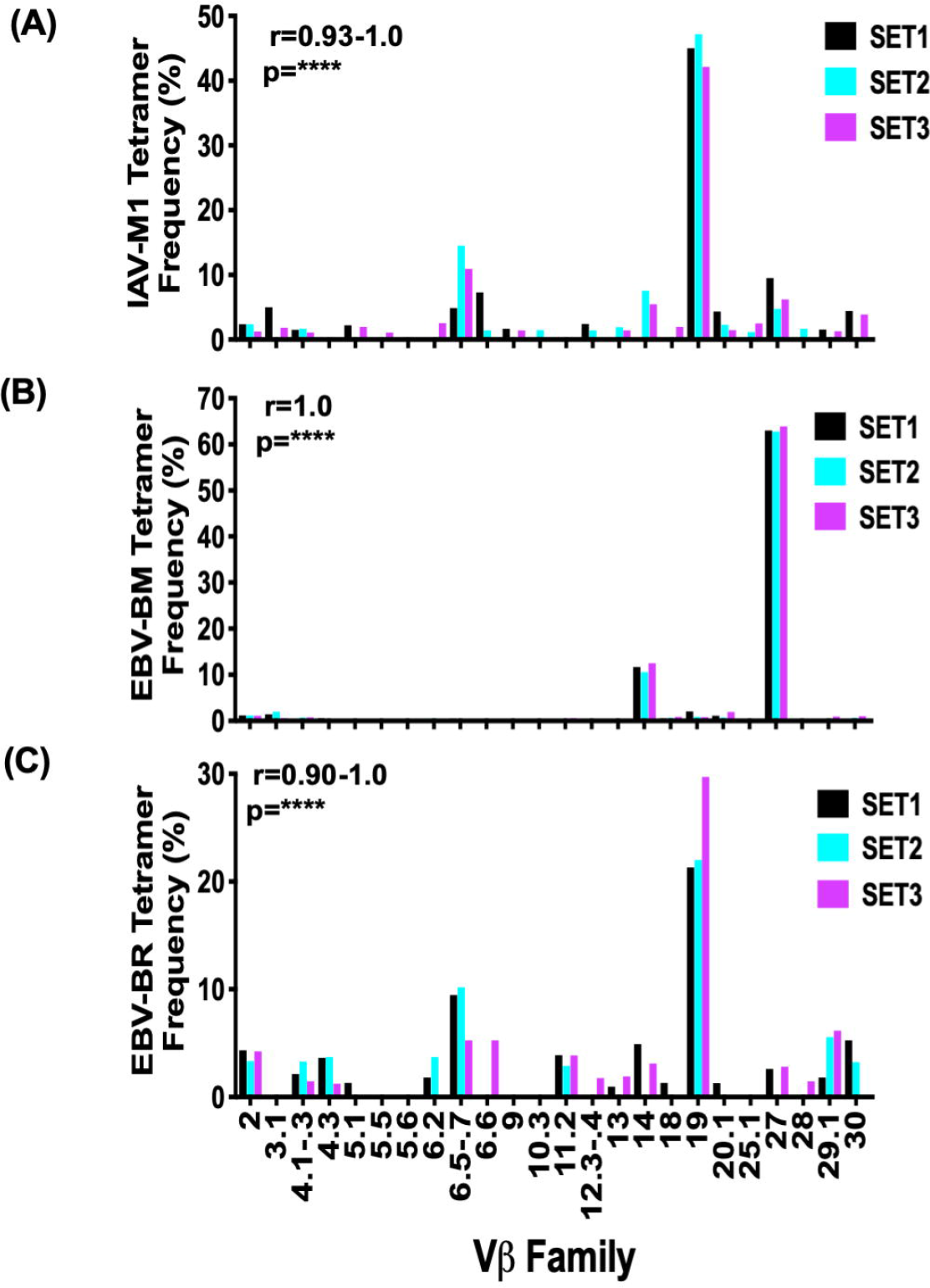
Highly reproducible TRBV repertoire monoclonal antibody staining for each epitope-specific response. Ex vivo sorted CD8 T cells freshly isolated from peripheral blood of donors were co-stained with (A) IAV-M1, (B) EBV-BM and (C) EBV-BR specific tetramers and TCR Vβ monoclonal antibodies. Triplicate staining was performed with two donors in parallel for all 3 epitope responses; representative bar graphs of donor D002 are shown. The Y-axis indicates the frequency TRBV family usage in the indicated tetramer positive populations. SET1 in black, SET2 in turquoise, SET3 in magenta. Triplicate staining frequencies were similar for each epitope. Pearson correlation was performed to compare all 3 data sets for each epitope response.

### Significant differences in epitope-specific TRBV usage in different age groups

Overall, there was a preferential usage of different BV families by each epitope-specific response for each age group (Figure 3A-C) and for each individual (Figure S2-S4). Specifically, the middle-age donors used significantly more BV28 in EBV-BM-specific responses than young donors (Figure 3B). Young donors (YSP) used significantly more BV3.1, BV5.1, BV6.2, and BV13 for EBV-BR responses compared to middle-aged (MSP) and older donors (OSP) (Figure 3C). There were clear differences in the hierarchy of BV usage for each epitope in each age group. The public TRBV19 was predominantly used by all age groups for IAV-M1 responses (Figure 3A). For EBV-BM-specific responses, the young donors had the most dominant usage of three BV families including the public BV9, BV20.1, and BV29.1, while middle-aged donors had the most dominant usage of the public BV20.1 and the less commonly used BV28 (Figure 3B). Interestingly, the OSP did not have any BV that were used more dominantly than others in the EBV-BM response, suggesting that each individual had a unique dominance profile and greater private specificity. For EBV-BR-specific responses which has a very diverse BV usage, there was dominant usage in YSP of BV5.1, BV6.2, BV13, and BV20.1, while there was no dominant BVs used in the MSP and OSP groups once again suggesting tremendous variability between donors or increased private specificity (Figure 3C). These data suggest that TRBV usage is changing with increasing age particularly for EBV-BM and EBV-BR epitope-specific responses.

**Figure 3.**
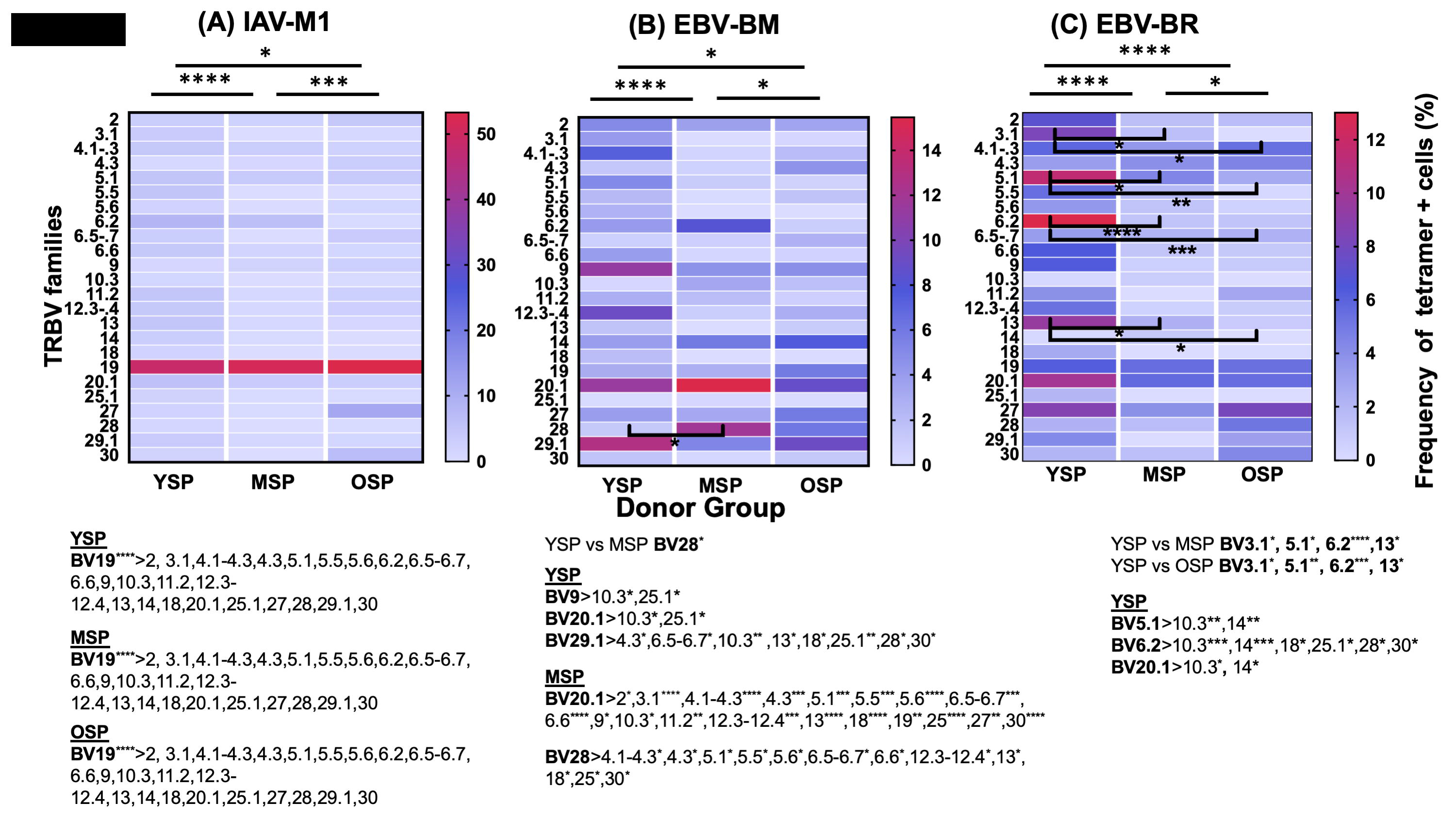
Preferential TRBV usage by each of the three major viral epitope-specific responses, IAV-M1, EBV-BM and EBV-BR, significantly differed between the different age groups. Ex vivo sorted CD8 T cells were co-stained with either (A) IAV-M1, (B) EBV-BM, or (C) EBV-BR-specific tetramers and TCR Vβ mAbs. Heatmap analyses show mean frequency used for each of the 24 TRBV families in the three different tetramer specific responses for each age group (YSP, n=14-23; MSP, n=19-20, OSP, n=9-10). The intensity of the color corresponds to the frequency of TRBV usage as shown in the right legend, light purple represents the lowest frequency, while red represents the highest frequency. Overall preferential TRBV usage differed for each epitope-specific response between all 3 age groups (Wilcox test with adjusted p value for multiple comparisons: * p<0.05, **p<0.01, ***p<0.001, ****p<0.0001). Preferential TRBV usage between age groups are indicated by black bars and significances (stars) on the heatmap. TRBV28 was used significantly more in middle-age than young donors for EBV-BM. Young donors used 4 TRBV, including BV3.1, BV5.1, BV6.2, and BV13, significantly more than middle-age or older donors. Below each epitope-specific heatmap is the hierarchical statistical analysis of which TRBVs were preferentially dominantly used by that epitope-specific response in each age group. This type of analyses showed BV19 was the most dominant TRBV in IAV-M1 specific responses in all age groups. For EBV-BM-specific responses, young donors had dominant usage of TRBV9, 20.1, and 29.1 at greater frequency than other TRBV; middle-age donors had dominant usage of BV20.1 and BV28 at greater frequency than other TRBV. For EBV-BR-specific responses, young donors had dominant usage of 4 TRBV families, including BV3.1, BV5.1, BV6.2, and BV13 more frequently than other families. Multivariant ANOVA with adjusted p values for multiple comparisons: * p<0.05, **p<0.01, ***p<0.001, ****p<0.0001. The TRBV data for each epitope for each individual is shown in pie charts in Figures S2-4.

### IAV-M1-specific BV usage correlated with age

Although all 3 age groups had a dominant usage of BV19 in the IAV-M1-specific response, older donors also used BV27 and BV30 (Figure 4A-C). We wanted to determine whether the use of BV19, 27, and 30 correlated with age. BV19 frequency did not directly correlate with age, but the analysis suggested that older donors may have a bimodal distribution of usage BV19 (and perhaps BV27) with some donors using predominantly BV19 family, while other older donors barely use it (Figure 4A, B). Interestingly, the frequency of BV27 and BV30 usage correlated directly with age (Figure 4C). These data suggest that some older donors are less dependent on BV19 family usage and develop a more diverse IAV-M1 specific TCR repertoire than other age groups, through the use of different BVs including BV27 and BV30 consistent with previously published data (69).

**Figure 4.**
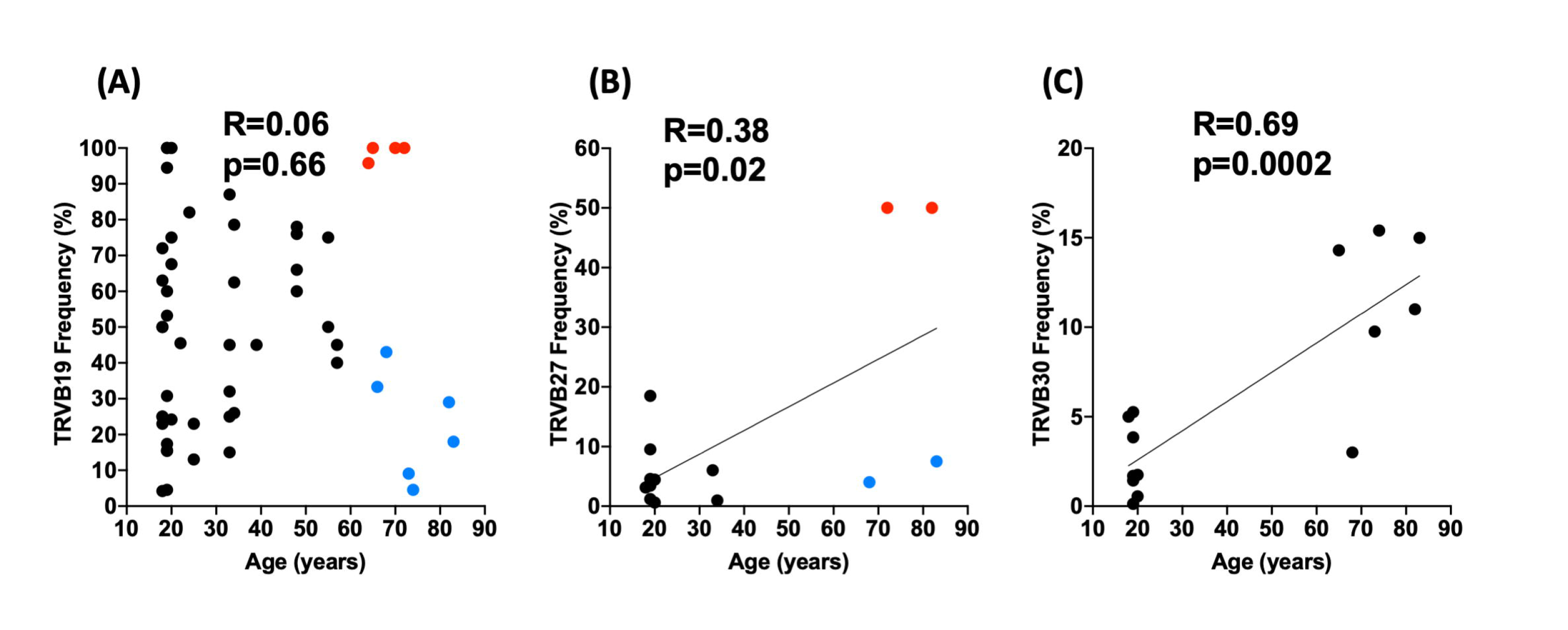
Changes in particular TRBV usage in IAV-M1-specific responses with increasing age. Ex vivo sorted CD8 T cells were co-stained with IAV-M1-specific tetramers and TCR Vβ mAbs. A) TRBV19 was predominantly used by young and middle age donors, while there is a bimodal distribution in older donors as indicated by the red and blue colored dots. B) Frequency of TBV27 usage correlated directly with increasing age although there was a somewhat bimodal distribution in the older donors. C) Frequency of TRBV30 usage correlated directly with increasing age. Spearman correlations shown.

### TRBV repertoire narrowing and potential cross-reactivity

If a particular TRBV is able to interact with more than one epitope this would suggest that it is more likely to be used by a crossreactive TCR. Therefore, we used the ability of particular TRBV in any age group to recognize more than one of the three epitopes as a measurable for potential crossreactivity. To further analyze the overall changes in TCR repertoire over age we did a Venn diagram analysis to examine how much overlap in dominant BV family usage was occurring between the three epitopes in the three different age groups as a measure of potential T cell cross-reactivity (Figure 5A-C). We found that young (8 BV) and older donors (3 BV) had more BVs shared amongst all 3 epitope-specific responses than middle age (1 BV) donors. Young donors had a broader TCR repertoire in which they used 5 different BV families specific to IAV- M1 only, while they shared 4 BVs between IAV-M1 and EBV-BR and one with EBV-BM. Interestingly, there were up to 8 BV families shared between all three epitope-specific responses (Figure 5A). In contrast, middle age donors had 6 shared BVs between EBV-BM and EBV-BR, but only one shared BV between all 3 epitope-specific responses (Figure 5B). Intriguingly, older donors had 4 BVs unique to the IAV-M1-specific response, 3 BVs unique to the EBV-BM-specific response and one BV family unique to the EBV-BR-specific response. There were 3 very dominant BVs shared by all three epitope-specific responses including the highly IAV-M1 public, BV19 and perhaps two unique public crossreactive TRBV, BV27 and BV4.3.

**Figure 5.**
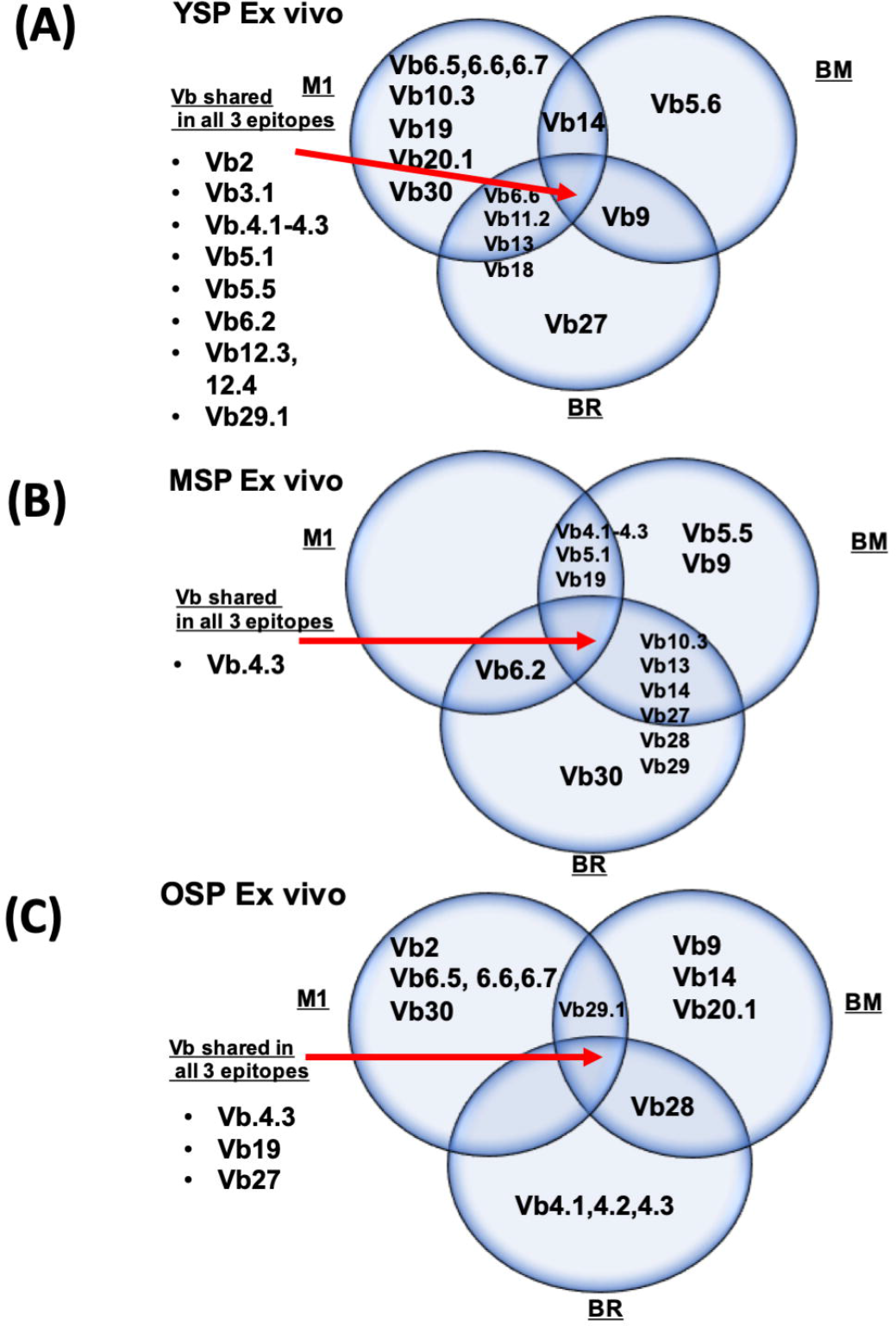
Significant changes in TCR repertoire with increasing age as assessed by overlap in TRBV usage by the 3 epitopes consistent with changes in cross-reactivity. Ex vivo sorted CD8 T cells from peripheral blood of young (YSP, n=20), middle-age (MSP, n=10), and older (OSP, n=10) donors were stained with IAV-M1, EBV-BM, and EBV-BR specific tetramers and TCR Vβ monoclonal antibodies. For each tetramer specificity, we analyzed only dominant TRBV usage in order to increase stringency (TRBV usage had to be ≥20% of the tetramer+ population to be included). The Venn diagram illustrates the unique (non-overlapping) and shared (overlapping) TRBV usage between the IAV-M1, EBV-BM, and EBV-BR epitope-specific responses of (A) YSP, (B) MSP and (C) OSP. Young donors had a total of 14, middle-aged had a total of 11 and older donors had a total of 5 BV out of the total 24 different BV tested (Chi square analysis, p=0.03) which overlapped between two or all three epitope-specific responses perhaps suggesting a narrowing and thus a preferential selection for particular cross-reactive TCR repertoire as individuals age

Young donors had a total of 14, middle-aged had a total of 11 and older donors had a total of 5 BV out of the total 24 different BV tested (Chi square analysis, p=0.03) which overlapped between two or all three epitope-specific responses perhaps suggesting a narrowing and thus a preferential selection for particular cross-reactive TCR repertoires as individuals age (Figure 5A- C). These results suggest that perhaps in younger donors the epitope-specific TCR repertoires were broader and therefore, more capable of cross-reactivity through multiple overlapping BV usage, while still maintaining unique BV for epitope-specific responses, whereas when individuals become older, the TCR repertoire narrowed as individuals become highly dependent on overlapping BV family usage. This is consistent with the concept of co-evolution as all three epitopes in the older donors are dominantly using the same three BV, BV19 (usually considered public for IAV-M1), BV27 and BV4.3 (which may be public for cross-reactive TCR between IAV- M1 and EBV-BM and or EBV-BR).

### TRBV repertoire to IAV-M1, EBV-BM and EBV-BR are continuously changing over 14 years enhanced by acute IAV infection in longitudinal studies

Most research performed in an effort to understand the impact of aging on CD8 memory T responses to viruses, has relied predominantly on cross-sectional studies (45–47). Using our sample repository, we performed a longitudinal study in which we tracked the IAV-M1, EBV-BM and EBV-BR specific CD8 T responses and TCR repertoire of 2 donors for over 14-15 years as they progressed from middle-age into the older age group (Figure 6,7). The TRBV repertoires to all three epitopes in both donors are continuously changing over the 14-15 years, although both donors appear to each have certain TRBV that show up as more dominant more frequently. These results are further supported by major fluctuations occurring in the epitope-specific TRBV repertoires of individuals in all of three age groups for donors that came in for more than one visit (Figure S2- S4). These findings are consistent with antigen-specific TCR repertoires continuously being modified overtime with increasing age, with some preference for particular dominant clonotypes. These two donors, D002 and D035 had documented cases of acute symptomatic IAV infection at least once during the 14-15-year period of these studies (Figure 6,7; Figure S5). Research has shown that the IAV-M1 frequency is relatively low, between 0.06-0.1%, when individuals do not have acute/active IAV infection (67, 70). IAV-M1-specific responses for both donors fluctuated, but increased by 3- to 6-fold during the acute IAV infections (Figure 6,7). Our data is consistent with previously published reports that showed an increase in the frequency of IAV-M1-specific CD8 T cells during acute IAV infection (71). Frequencies for EBV-BM and EBV-BR-specific cells remained relatively stable for each donor before acute IAV (Figure 6,7). However, during the acute IAV infection EBV-BR-specific responses also increased 2 to 5-fold in these donors (Figure 6,7), while EBV-BM-specific responses increased 3-fold in donor D002 going from 2% to 6%. This could result from reactivation of EBV, but we did not find a measurable increase in EBV genome copies in the B cells of these two donors at the time of the increase in EBV-specific CD8 T cells.

**Figure 6.**
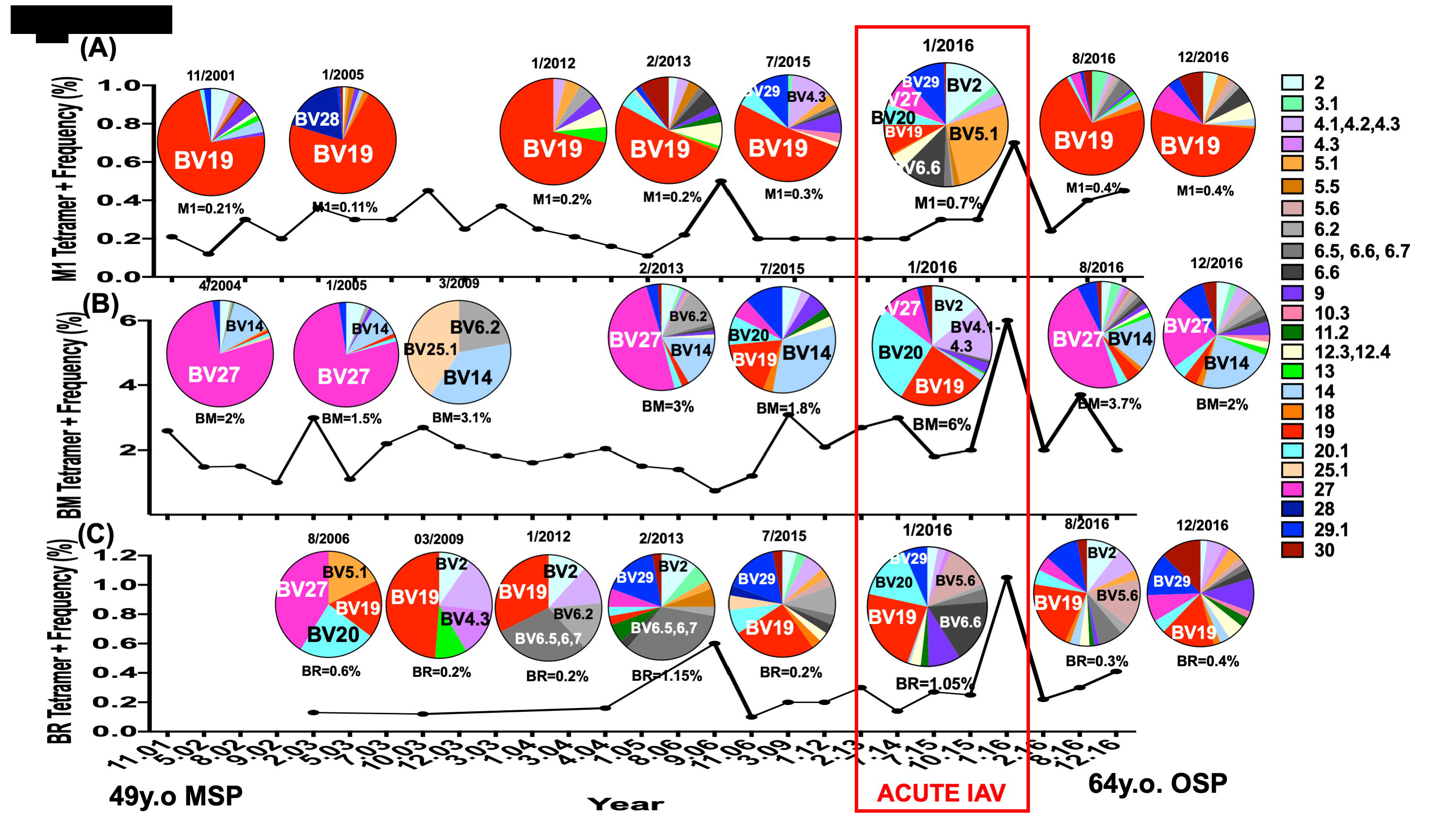
IAV-M1 and EBV-BM, EBV-BR TCRBV repertoires co-evolve longitudinally overtime in donor D002 from middle-age to elderly enhanced by acute IAV infection. Ex vivo sorted CD8 T cells were co-stained with either (A) IAV-M1, (B) EBV-BM or (C) EBV-BR specific tetramers and TCR Vβ mAbs. Tetramer-specific CD8 T cell frequencies and TCRBV repertoire were examined and recorded over 15 years, from 11/2001 to 12/2016. During 1/2016, the donor (D002) became ill with acute symptomatic influenza A, for which point, the corresponding CD8 T cell frequencies and TCRBV repertoires were highlighted by a red box. TRBV repertoire are presented in the form of pie charts. The line graph illustrates the frequencies of freshly isolated tetramer+ CD8 T cell populations at several time points for donor, D002. The IAV-M1 tetramer+ frequency was relatively stable (0.1-0.2%) until the acute IAV infection in 1/2016, when it increased to 0.7%. At that same time point, the EBV-BM and EBV-BR responses also increased to an all-time high of 6% and 1%, respectively. In the IAV-M1 specific response, TRBV19 dominated the response until the acute IAV infection, when its frequency dramatically decreased, while the frequencies of other TRBV that overlapped with public EBV-BM or EBV-BR BV usage increased, including TRBV20, TRBV2, BV5.1, BV6.6, BV27 and BV29. The TRBV repertoire for EBV-BM changed over the years but there was some consistent usage of either BV27 or BV14 dominating. However, during the acute IAV infection the frequency of BV19 usage increased (usually dominant in IAV-M1-specific responses), as well as the usage of BV20.1, BV4.1,4.2,4.3 family, and BV2, while the usage of BV27 and BV14 dramatically decreased. EBV-BR TRBV repertoire changed continuously over the years, with TRBV19 usage being one the most consistent findings including during acute IAV. However, during acute IAV infection there was increased usage of BV20, which was also present in the EBV-BM and IAV-M1 responses. The date and tetramer-specific frequency of each response are shown above or below the TCR repertoire pie chart.

**Figure 7.**
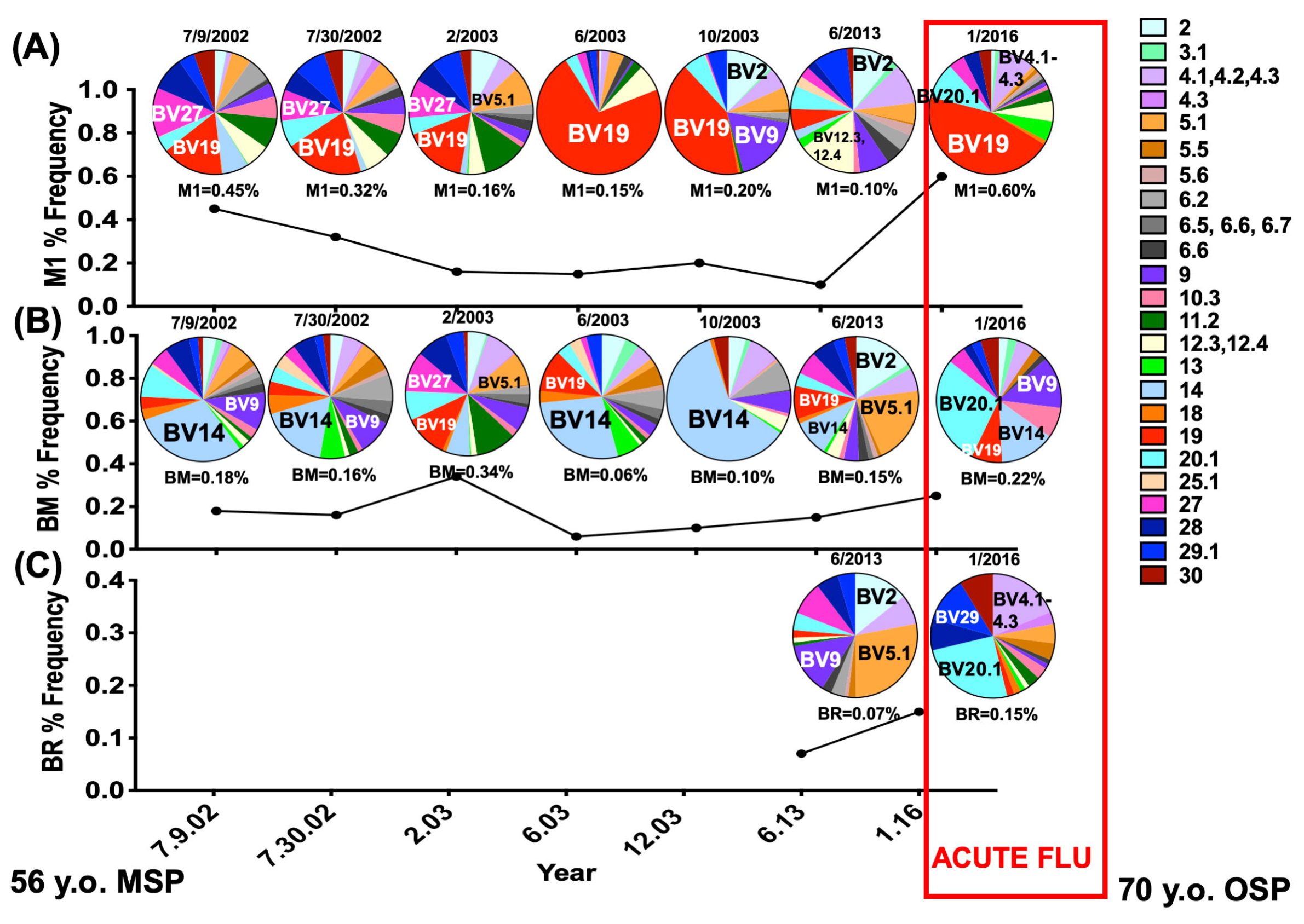
IAV-M1 and EBV-BM, EBV-BR TRBV repertoires co-evolve longitudinally in Donor D035 overtime from middle-age to elderly enhanced by acute IAV infection. Ex vivo sorted CD8 T cells were co-stained with either (A) IAV-M1, (B) EBV-BM or (C) EBV-BR specific tetramers and TCR Vβ mAbs. Tetramer-specific CD8 T cell frequencies and TRBV repertoire were examined and recorded over 14 years, from 11/2002 to 12/2016. During 1/2016, the donor (D035) became ill with acute symptomatic IAV infection, for which point, the corresponding CD8 T cell frequencies and TRBV repertoires are highlighted by a red box. The line graph illustrates the frequencies of freshly isolated tetramer+ CD8 T cell populations at several time points for donor, D035. The IAV-M1 tetramer+ frequency was relatively stable from 2003 to 2013 (0.1- 0.2%) until the acute IAV infection in 1/2016, when it increased from 0.15 in the previous visit to 0.6%. At that same time point, the EBV-BM frequency was stable (0.1-0.3), while the EBV-BR responses increased from 0.07% to 0.15% during acute IAV. In the IAV-M1 specific response, the TRBV changed over the years with TRBV19 becoming more dominant between 2002 and 2003 when it dominated the response, but by 2013 it was no longer dominant. Then during the acute IAV infection when TRBV19 frequency dramatically increased, while the frequencies of other TRBV that overlapped with EBV-BM or EBV-BR responses were still present including TRBV20, and BV4 family. The TRBV repertoire for EBV-BM changed over the years but there was some relatively consistent usage of BV14. However, during the acute IAV infection the frequency of BV20.1 usage increased in both EBV-BM and EBV-BR responses, (also dominant in IAV-M1- specific responses), as well as the usage of BV9, and BV14 usage decreased. EBV-BR TRBV repertoire data was only available in 2013 and 2016 largely because his EBV-BR tetramer+ cells were poorly detectable by tetramer. However, during acute IAV infection there was increased usage of BV20.1, which was also present in the EBV-BM and IAV-M1 responses and of BV4 family which was also dominant in the IAV-M1 tetramer+ cells. The date and tetramer-specific frequency of each response are shown above or below the TCR repertoire pie chart.

In D002 we found new double tetramer positive cells, when co-staining with either IAV- M1 and EBV-BM or EBV-BR specific tetramers (Figure S5), which suggests that acute IAV infection could increase the frequency of both IAV- and EBV-specific and cross-reactive CD8 memory T cells (Figure 6,7). The frequency of BV19 usage increased in donor D002 in EBV-BM and EBV-BR specific TRBV repertoires, despite it having been established as a public response to IAV-M1, while it decreased in the IAV-M1 response (Figure 6). Furthermore, in D002 the IAV- M1 response had dominant usage of BV2, BV29, BV27 and BV20 (72) all public BV for EBV- BM responses, while the dramatically increased usage of BV5.1 and BV6.6 are BV that are usually associated with EBV-BR responses (72). The TRBV repertoire for EBV-BM specific responses maintained a dominant use of TRBV14 for both donors until acute IAV infection, during which BV20 usage increased in both EBV-BM and -BR specific responses, as well as IAV-M1 responses (Figure 6,7). This would potentially be consistent with increased TCR cross-reactivity between EBV-BM and EBV-BR specific responses as well as IAV-M1, which we have previously reported in AIM donors(67).

In one of the donors, D002, we have two post-acute IAV visits, 8 and 12 months after the infection, where the TRBV repertoire at least in IAV-M1 and EBV-BM appears to be shifting back to a repertoire more similar to before infection, although far from identical. We assume the presence of the persistent EBV, which continuously reactivates (in the absence of IAV) is driving these changes, but until we look at the single cell clonotype level we do not know whether these are same or new clonotypes with similar features. This would be consistent with the observation that these TCR repertoires are co-evolving. It may also suggest that the dominance of BV19 usage in the IAV-M1 response is actually driven by its crossreactivity with EBV-BM or EBV-BR. Interestingly, BV19 usage does not become dominant in the IAV-M1 response until age 16 when many individuals in Western Society become infected with EBV, while almost 100% have had IAV by age 3 (73, 74). Unfortunately, we did not have PBMC from these two donors when they were in the young age group (YSP) or even before the age of 40 years old as many of the greatest differences in diversity and TRBV usage occur between the YSP and the other two age groups. However, these data suggest that epitope-specific CD8 TCR repertoires for these three epitopes are continuously changing and that acute IAV infection may contribute to these changes by increasing the selection for cross-reactive TCR between the 3 epitopes.

## Discussion

Here we show in cross-sectional studies that IAV-M1, EBV-BM and EBV-BR specific CD8 TCR repertoires are different in young, middle-aged and older donors with a narrowing of their TRBV family usage with increasing age to different degrees for each epitope. In fact, narrowing of EBV-BM and EBV-BR-specific BV usage correlated with increasing age. Although IAV-M1-specific TRBV repertoire narrowed from young donors to middle age donors, it did not directly correlate with increasing age, but there was clear evidence that the BV usage was changing with age. The dominant IAV-M1 BV19 family usage appeared to become bimodal in the older age group and interestingly BV30 usage did directly correlate with age. For the EBV epitope-specific responses there was preferential usage of particular TRBV and changes in the hierarchy of BV usage in the different age groups. Furthermore, longitudinal studies suggest that when cross-reactive responses are known to be present, as is the case with the three epitopes we studied, they may influence the evolution of virus-specific repertoires particularly during acute viral infection. We were fortunate to be able to track TCR repertoire changes in 2 donors longitudinally over 14- 15 years (middle age to older) showing that there were changes in the TCR repertoire of IAV-M1, EBV-BM and EBV-BR-specific responses over time. These two donors experienced acute symptomatic IAV infection and there was evidence these repertoire alterations may be influenced by TCR cross-reactivity, which is enhanced during acute IAV infection. These studies show that virus-specific TCR repertoires change as an individual ages leading to narrowing of the repertoire consistent with co-evolution in the presence of CD8 T cell cross-reactivity.

We were able to use 3 different age groups in our cross-sectional study to show that changes were occurring in TRBV usage overtime including the narrowing of the repertoires with increasing age. The identification of unique BVs used by each age group and between age groups for all 3 epitopes, highlighted the changes that occur with increasing age. In addition, we showed unique data using an under-studied young age group, 18-19-year old healthy EBV-serologically positive donors, where antigen-specific TCR repertoires appear to still have increased diversity, which only begins to narrow in middle age, the group that is most often studied for TCR repertoire analyses. We determined the IAV-M1, EBV-BM and EBV-BR specific TRBV diversity of young adults and their preferential BV family usage hierarchy for all 3 epitopes. We show data from the type of longitudinal studies that are rarely performed or published, in which we follow 2 individuals over 14 years, from middle-age to older, tracking their antigen-specific responses and TRBV repertoire to two common human viruses using their 3 most well-conserved, well-characterized, and immunodominant epitope responses. These studies were important in helping us to identify how these virus-specific responses co-evolve, especially during an acute infection. We found that an acute IAV infection enhances TCR cross-reactivity among all 3 epitopes responses as suggested by the fact that all 3 epitope specific responses increased during infection and they began to share well-characterized, public BV families like BV20 (usually public for EBV-BM), BV19 (usually public for IAV-M1) and BV27 (maybe public for EBV and IAV T cell crossreactivity).

In cross-sectional studies for all 3 epitope-specific responses there was evidence of narrowing of TRBV repertoire usage with increasing age (Figure 1) and dominant usage of 3 overlapping BV families by all 3 epitopes, BV19, BV27 and BV4.3 (Figure 5). These data could be consistent with increased dependence on narrowed, cross-reactive responses (Figure 5). In parallel, IAV-M1 repertoire appeared to change with age, as it had a narrower BV usage than EBV-BM and EBV-BR in young and middle-aged donors (Figure 1). The IAV-M1-specific repertoire also had a highly focused TRBV family usage, BV19, which became broader in some older donors. The IAV-M1-specific response may have the most narrowed TRBV repertoire diversity, because this epitope-specific response is predominantly selected based on an interaction between the TRBV chain and peptide/MHC complex (56, 75), while in EBV-BM and EBV-BR responses the TRAV chain is known to play an important role (61, 64, 76). In addition, these results are consistent with our previously published data and that of others, which show BV19 as the most dominant and public response to IAV-M1 in HLA-A2.01+ individuals (56, 77, 78). One previous study has also shown that a portion of older donors tend to use less BV19 with age, instead using BV27 among others, which is consistent with our TRBV repertoire mAb staining results (79). This drifting away from the usage of the BV19-expressing TCR specific to IAV-M1, may increase susceptibility to IAV infection in the older age group. EBV-BM, on the other hand, has several published public TCRs, including TRBV20, 29, 14, 9 and 2 (61–63, 80). While few papers have described the public BVs for EBV-BR, our previous work has suggested that EBV-BR is predominantly selected on peptide/MHC interaction with the TCRα chain and therefore appears to be more flexible in TRBV repertoire usage, leading to identifying often 4-6 dominant BVs in one donors and very different BV family usage between donors (61, 64).

Many studies have shown that TCR memory and naïve phenotype T cell repertoires narrow with increasing age most likely as a result of thymic involution and gradual loss of the remaining T cells (45, 46). Most previous research did not examine BV family preference by age groups, but instead, they explored preferential usage of certain BVs for particular epitopes in pathogens including, IAV, EBV, CMV, HCV and HIV often studying middle age donors (56, 61, 63, 64, 79, 81–83). Our cross-sectional studies which showed for all 3 epitopes that TRBV usage evolved with age with different preferential usage of particular BV by each age group and between age groups would suggest that this is not a random event. It might not be surprising that young donors, who have broader overall TCR repertoires would use more BVs for all 3 epitope responses than the other age groups and potentially have a much broader potential for cross-reactive responses as suggested by the number of overlapping BV families between the epitopes. This might be part of the explanation for why young individuals in their early twenties are more prone to developing autoimmune responses. But what is the mechanism driving preferential usage of particular BV families by a certain age group for these two viruses? It is possible that recurrent stimulation with EBV lytic epitopes every time the virus goes into low level lytic cycle or recurrent infection with IAV over the years leads to the selection of certain better fit or higher affinity clones. However, if that was the case, you would expect that older donors would have the best protection against both viruses and yet they do not. It is of course possible, that the high affinity clones become exhausted and disappear as individuals age but that has usually been only described with very high sustained viral loads (84, 85).

Our data would suggest that in fact, TCR cross-reactivity could be another mechanism that contributes to this evolution of TCR repertoire as evidenced by our Venn diagram analyses of the cross-sectional data as well as the significant changes over time and particularly during acute IAV infection in our longitudinal studies of 2 donors. If a particular TRBV is able to interact with more than one epitope this would suggest that it is more likely to be used by a crossreactive TCR. Therefore, in these studies we used the ability of particular TRBV in any age group to recognize more than one of the three epitopes as a measurable for potential crossreactivity. The cross-sectional study allowed us to demonstrate the *potential* for *ex vivo* IAV-and EBV-specific CD8 TCR cross-reactivity with increasing age. In a rudimentary assessment of potential cross-reactivity, we examined the overlap of TRBV usage between *ex vivo* epitope-specific responses in YSP, MSP, and OSP using a simple Venn Diagram. We found that YSP maintained TRBV usage that was both unique to each epitope-specific response and common between all 3 epitope-specific responses. These results were consistent with our TCR diversity data, which showed that YSP have great IAV-M1, EBV-BM, and EBV-BR CD8 TCR repertoire diversity compared to MSP and OSP (Figure 1). These data are also consistent with literature that finds that the overall CD8 TCR diversity for younger individuals are greater than older individuals (35, 45–47, 69, 86). However, the middle-age donors only shared 1 TRBV, TRBV4.3, amongst all 3 epitope-specific responses. MSP shared several TRBV between EBV-BM and EBV-BR specific responses, as well as, between IAV-M1- and EBV-BM-specific responses. These results were also consistent with our antigen-specific CD8 TCR diversity data, which shows a significant decrease in IAV-and EBV-specific CD8 TCR diversity with increasing age. Young donors had a total of 14, middle-aged had a total of 11 and older donors had a total of 5 BV out of the total 24 different BV tested (Chi square analysis, p=0.03) which overlapped between two or all three epitope-specific responses perhaps suggesting a narrowing and thus a preferential selection for particular cross-reactive TCR repertoire as individuals age (Figure 5A-C). These results suggest that perhaps in younger donors the epitope-specific TCR repertoires were broader and therefore, more capable of cross-reactivity through multiple overlapping BV usage, while still maintaining unique BV for epitope-specific responses, whereas when individuals become older, the TCR repertoire narrowed as individuals become highly dependent on overlapping BV family usage. This is consistent with the concept of co-evolution as all three epitopes in the older donors are dominantly using the same three BV, BV19 (usually considered public for IAV-M1), BV27 and BV4.3 (which may be public for cross-reactive TCR between IAV-M1 and EBV-BM and or EBV-BR) (58, 80, 87–96).

The longitudinal data, in which we followed 2 donors from middle-age to older for over 14-15 years, allowed us to make 3 major unexpected discoveries that demonstrated potential TCR cross-reactivity. We discovered that during acute influenza infection: all 3 epitope specific responses increased, all 3 epitope-specific TCR repertoires changed, and that in at least 1 donor, TRBV19 usage increased in all 3 epitope specific repertoires. TRBV19 is a well published public response to IAV-M1 (58, 69, 87, 94, 96–99). The increase in the presence of TRBV19 suggests that TRBV19 has features in its CDR3 loop that would allow it to recognize both EBV-BM, and EBV-BR. Another indication of TCR cross-reactivity during acute influenza infection was the increased usage of TRBV20 in both EBV-BM and –BR-specific responses, despite TRBV20 having been identified as a public response to EBV-BM (61, 88–90, 92, 93, 95, 100–103). These data suggest co-evolution of the TCR repertoire, in which, public features of the IAV- and EBV-specific repertoires (TRBV19 and TRBV20) are being shared amongst the 3 epitope specific responses, especially during acute IAV infection. In the absence of TCR repertoire co-evolution, we would not expect there to be sharing of public BVs amongst IAV-and-EBV-specific responses, particularly during an acute infection, in which there is an inflammatory environment. As we have previously identified TCR cross-reactivity in healthy donors, these data also suggested that acute influenza infection may enhance TCR cross-reactivity (80, 104). Overall, this longitudinal study provided a snapshot of what happens to IAV-and EBV-specific responses and TCR repertoire during a co-infection with IAV and EBV, providing a model for two unrelated viruses that cause acute and persistent viral infections, and demonstrated how an acute infection may potentially increase cross-reactive responses as well as perturbations to the TCR repertoire.

Our data would suggest that in fact, TCR cross-reactivity could be a mechanism that contributes to this evolution of TCR repertoire as evidenced by significant changes over time and particularly during acute IAV infection in our longitudinal studies of 2 donors. In particular, IAV- M1 and EBV-BM and EBV-BR epitope-specific CD8 T cell tetramer frequency increased and changes in TRBV usage occurred in the two donors who experienced acute symptomatic IAV infection. We also demonstrated increased frequency of IAV- and EBV-specific double tetramer positive cells during acute IAV infection. Studies in EBV infection and recently in the COVID19 pandemic have shown that cross-reactive responses can become activated during acute viral infections and may play a role in a broad range of symptoms from severe immunopathology to protection (99, 105, 106). For instance, we have shown that not only the frequency of IAV-M1- specific CD8 T cells correlates with the severity of AIM but the particular TRBV usage of the reactivated IAV-M1-specific cells (106). In direct contrast, we have shown that particular unique IAV-M1 TCR cross-reactive with EBV-BM and EBV-BR may play a role in protection from EBV seroconversion in those 5% of individuals that remain EBV serologically negative past their 40’s (99). Mouse studies, but few if any human studies, have shown that the presence of a silent persistent infection can impact the CD8 T cell responses and TCR repertoire during an acute new viral infection, creating an environment conducive to TCR cross-reactivity as both viruses are present at the same time (107–109).

The narrowing of the TCR repertoire diversity in older donors may result in increased use and retention of cross-reactive TCRs as the CD8 T cell pool gradual declines with increasing age. Older donors may select cross-reactive TCRs that can respond to both IAV and EBV. More studies are needed to understand the impact that these particular BVs may have on controlling both the IAV and EBV infections. Many researchers are trying to use TCR repertoire analyses to identify novel epitopes in diseases, infections and cancer, to find what role TCR plays in disease severity, and to identify the role of cross-reactivity in TCR repertoire diversity (110–112). Therapeutically increasing the TCR repertoire diversity to include particular BV families may decrease susceptibility to infections and cancer but could enhance autoimmunity. One day we may be able to harness the power of TCR repertoire to therapeutically engineered T cells in order to treat cancers and severe viral infections, while at the same time decreasing the risk of autoimmunity. These studies are significant in providing understanding of how CD8 memory T cell responses and TCR repertoire diversity change with increasing age in the presence of two viruses that are likely present together in individuals at a given time. IAV and EBV infections have significant consequences on quality of life and the healthcare system. IAV costs the United States health care system and economy an average of $11.2 billion, while EBV-attributed cancers lead to nearly 8 million deaths globally (113, 114). As the life expectancy of individuals in the United States becomes higher than at any time in history, at 78.6 years, it will become increasing critical to advance our knowledge and understanding of how the immune system changes with age to decrease susceptibility to infection, mortality, and health care burdens (115). These types of studies will also be important in understanding the role TCR repertoire plays in response to viral infections.

## Materials and Methods

### Study population

This project used donors from the University of Massachusetts Amherst (UMA) EBV Serological Surveillance Cohort. Donors were healthy, chronically infected EBV seropositive, young adults enrolled at University of Massachusetts Amherst (UMA). Donors were recruited and enrolled into the clinic, as freshman (first year) students for first year orientation and were followed until their senior year (last year) of college. Blood was drawn from these donors once per semester or twice a year. Our lab acquired up to 30ml of peripheral blood from HLA-A2.01 donors. The donors were on average 18.8 years old. University of Massachusetts Medical School (UMMS) volunteers are HLA-A2.01 donors that are middle-age (25-59 years old) and older (60-93 years old). IAV and EBV-specific immunity was confirmed using antigen specific tetramers. EBV serology was confirmed through the detection of IgG specific antibodies for the viral capsid protein. These donors were allowed to donate up to 150ml of blood in 3 months (IRB). This study was approved by the Institutional Review Board (IRB) committee at University of Massachusetts Medical School, Worcester, Massachusetts. All donors in this study provided informed consent.

### HLA-Typing

Monoclonal antibody specific to HLA-A2 (clone BB7.2, BioLegend, San Diego, CA, was added to 100ul of whole blood, incubated for 30 minutes at room temperature (RT) and washed with 1ml of Hank’s Balanced Salt Solution (HBSS) (Gibco, Grand Island, NY). Cells were spun at 1330rpm for 4mins, at RT. Red blood cells were lysed using 2ml of 1X BD FACS Lysing Solution (Becton Dickinson, Waltham, MA) and incubation for 10 minutes at 37°C. Cells were washed and re-suspended in 300ul FACS buffer (500ml HBSS, 2% Fetal Calf Serum) and analyzed on the LSRII (Becton Dickinson, Waltham, MA).

### PBMC Isolation

A 1:1 dilution was used when combining fresh whole blood and Hank’s Balanced Salt Solution (Gibco, Grand Island, NY). Approximately 25mls of this mixture was layered over 15mls of Ficoll-Paque Plus (GE Healthcare Bio-Sciences, Pittsburgh, PA). The cells were spun at 1800rpm for 40 minutes with no brake at RT. The buffy coat containing the PBMC was taken carefully and washed twice with 20ml HBSS.

### CD8 T cell isolation

The PBMC were counted and re-suspended in 20μl of anti-CD8 micro-beads (Miltenyi Biotech, Auburn, CA) and 80μl of MACS buffer [4°C Phosphate-buffered saline, 2.5g of Bovine Serum Albumin (Sigma-Aldrich, St.Louis, MO), 2ml 0.5M EDTA [pH 8.0] (Invitrogen, Grand Island, NY) degassed with sterile mesh filter] per 10^7^ cells. The PBMC were incubated for 15 minutes in the dark at 4°C and then washed with 20ml of MACS buffer. The CD8 T cells were isolated using Miltenyi Biotech MACS system as previously described(65).

### Tetramers and Dextramers

The IAV-M1 tetramer, EBV-BMLF1 tetramer, EBV-BRLF1 tetramer (NIH Tetramer Core Facility, Atlanta, GA), and IAV-M1 dextramer (Immudex, Copenhagen, Denmark) were assembled using the sequences above and conjugated to either allophycocyanin (APC) or phycoerythrin (PE). Tyrosinase (NIH Tetramer Core Facility, Atlanta, GA) and CMV_pp65_ (our tetramer core facility and NIH Tetramer Core Facility, Atlanta, GA) were used as negative controls for all experiments.

### Extracellular Staining

For staining, 3 x 10^5^ freshly isolated cells were placed into each well of a 96-well plate. Cells were co-stained with tetramers or dextramers and activation surface markers for 30 minutes at RT. Cells were washed twice with FACS buffer (500ml Hank’s Balanced Salt Solution with 2% Fetal Calf Serum) and fixed using 100μl of Cytofix (Becton Dickinson Biosciences, San Jose, CA) for 5 mins. in the dark at room temperature. The cells were washed with and re-suspended in FACS buffer and analyzed on the LSRII (Becton Dickinson, Waltham, MA). All activation surface marker antibodies were purchased from BioLegend (San Diego, CA): CTLA-4 (CD152) PE-Cy7(clone: BNI3), PD-1(CD279) APC-Cy7 (clone: NAT105), HLA-DR PE (clone: L243), KLRG1 BV421 (clone: SA231A2), CD69 Alexa Fluor 700 (clone: FN50), 2B4 (CD244) FITC (clone: C1.7).

### TCR V beta repertoire staining

The TCR V beta repertoire kit contained antibodies to over 76% of V βeta families (Beckman Coulter, Fullerton, CA). The cells were stained with these antibodies in combination with tetramers or dextramers for 20 minutes at RT to determine the TRBV repertoire of antigen specific cells. The cells were washed in FACS Buffer and placed at 4°C.

### Statistics

2 way-ANOVA multi-variant analysis with adjusted p values for multiple comparisons was applied to analyze multiple donor groups. Simpson Diversity Index was used to analyze TCR diversity, Pearson’s and Spearman’s correlation coefficients were used for analyses of correlations.

## Acknowledgements

We would like to thank Dr. Eva Szomolanyi-Tsuda for here assistance in reading and editing our manuscript. We would also like to thank Dr. George Corey, Jessica Conrad, Robin Brody and Margaret McNamara for their assistance with obtaining donor samples for our studies.

## Supplemental figure legends

Supplemental Table 1. Characteristics of the study populations.

**Figure S1. Ex-vivo EBV-BM frequencies significantly increases in middle-aged adults.** *Ex vivo* tetramer staining on magnetically sorted CD8 T cells of an IAV-immune and EBV seropositive young, middle-age, and older donor that are HLA-A2+. We tetramer stained for immunodominant IAV-M1 and EBV-BM and EBV-BR epitopes and tyrosinase, a self-epitope on melanocytes as negative control. The graph illustrates the frequency of CD8 tetramer positive T cells of EBV seropositive young (n=20; shown as dark blue circle), middle aged (n=10; shown as red square), and older donors (n=10; shown as light blue triangle). The frequencies for EBV-BM tetramer+ cells are significantly higher in middle-aged donors than in either young or older donors. Multivariant ANOVA with adjusted p values for multiple comparisons: **p<0.01, ***p<0.001.

**Figure S2. Private specificity of IAV-M1 specific TRBV repertoires.** Sorted CD8 T cells freshly isolated from peripheral blood of young (n=23), middle-age (n=19), older (n=10) donors were stained with IAV-M1 specific tetramer and a TCR Vβ repertoire mAb kit over several visits. TRBV repertoires are presented in the form of pie charts. The individual TRBV pie charts are followed by a union TRBV pie chart that compiled all donors in each age group, which allowed us to present an overall summary of BV usage in each age group. The time between visits for young donors was 6 months to one year. Most middle-age and older donors were followed over a year, while two donors were tracked up to 14 years, from middle-age to older, including D002, and D035. Since not only was there significant variability in TRBV usage between donors (private specificity) but also within in each donor over multiple visits, each visit was considered a unique data point.

**Figure S3. Private specificity of EBV-BM specific TRBV repertoires.** Sorted CD8 T cells freshly isolated from peripheral blood of young (n=15), middle-age (n=18), older (n=9) donors were stained with EBV-BM specific tetramer and a TCR Vβ repertoire mAb kit over several visits. TRBV repertoires are presented in the form of pie charts. The individual TRBV pie charts are followed by a union TRBV pie chart that compiled all donors in each age group, which allowed us to present an overall summary of BV usage in each age group. The time between visits for young donors was 6 months to one year. Most middle-age and older donors were followed over a year, while 2 donors were tracked up to 14 years, from middle-age to older, including D002, and D035. Since not only was there significant variability in TRBV usage between donors (private specificity) but also within in each donor over multiple visits, each visit was considered a unique data point.

**Figure S4. Private specificity of EBV-BR specific TRBV repertoires.** Sorted CD8 T cells freshly isolated from peripheral blood of young (n=19), middle-age (n=11), older (n=9) donors were stained with EBV-BR specific tetramer and a TCR Vβ repertoire mAb kit over several visits. TRBV repertoires are presented in the form of pie charts. The individual TRBV pie charts are followed by a union TRBV pie chart that compiled all donors in each age group, which allowed us to present an overall summary of BV usage in each age group. The time between visits for young donors was 6 months to one year. Most middle-age and older donors were followed over a year, while two donors were tracked up to 14 years, from middle-age to older, including D002 and D035. Since not only was there significant variability in TRBV usage between donors (private specificity) but also within in each donor over multiple visits, each visit was considered a unique data point.

**Figure S5. Increased ex vivo frequency of virus-specific IAV-M1, EBV-BM, and EBV-BR, and cross-reactive IAV-M1+EBV-BM, IAV-M1+EBV-BR tetramer+ cells during acute IAV infection**. FACS plots show the results of freshly isolated sorted CD8 T cells from the peripheral blood of donor D002 that were stained ex vivo with either IAV-M1, EBV-BM, or EBV-BR specific tetramers or co-stained with two of these tetramers one year before acute IAV infection in 7/2015 (A) and subsequently during acute symptomatic IAV infection in 1/2016. New populations of double-tetramer positive cells (IAV-M1+IAV-BM and IAV-M1+EBV-BR) appeared during acute IAV.

