## Supplementary figures and images for "Co-evolution of human influenza A and Epstein Barr virus-specific CD8 ex vivo memory T cell receptor BV repertoires with increasing age"

### Figure S1

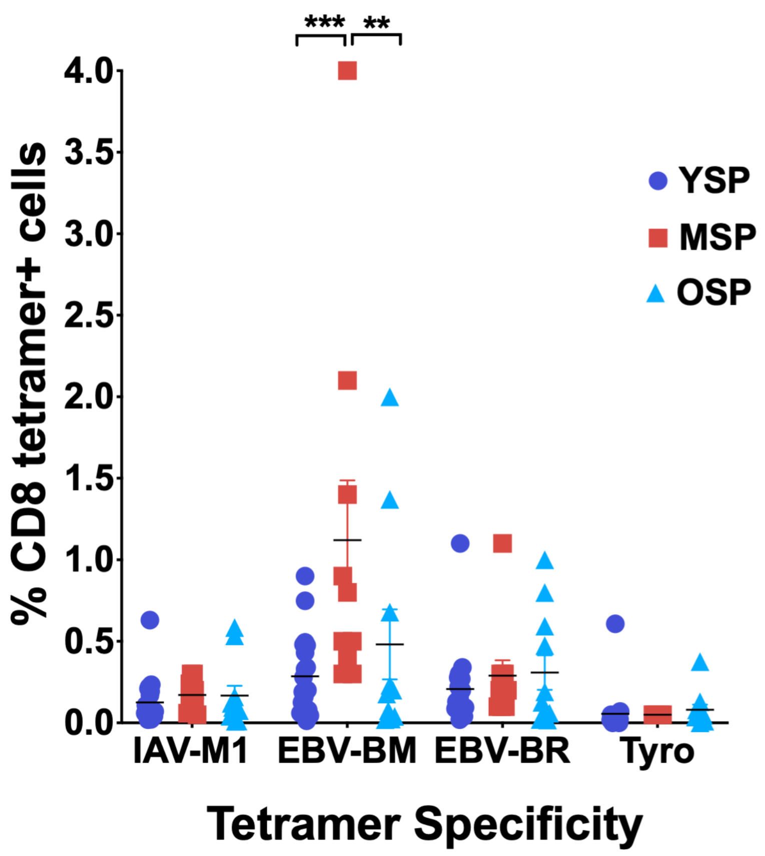

### Figure S2

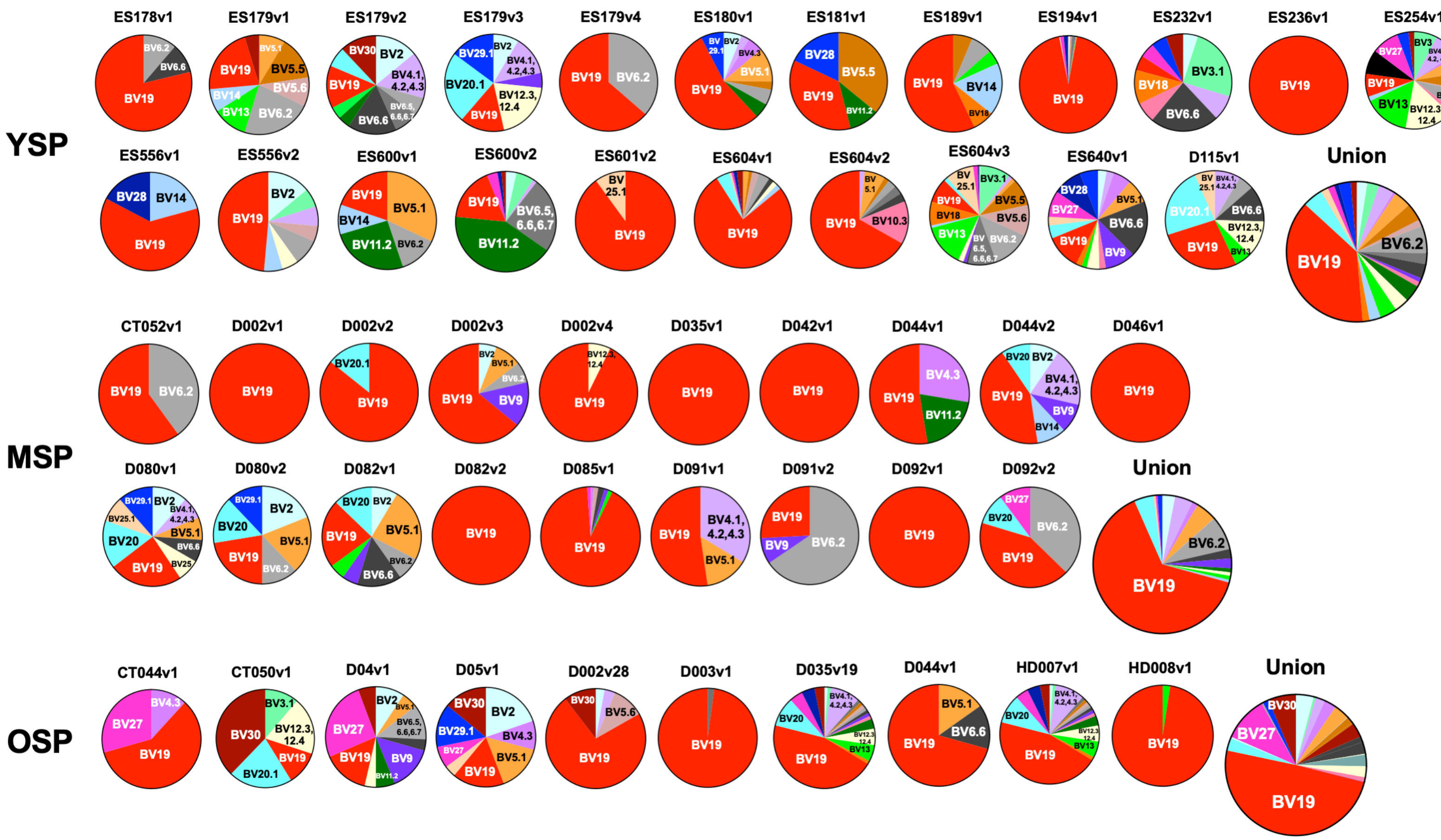

### Figure S3

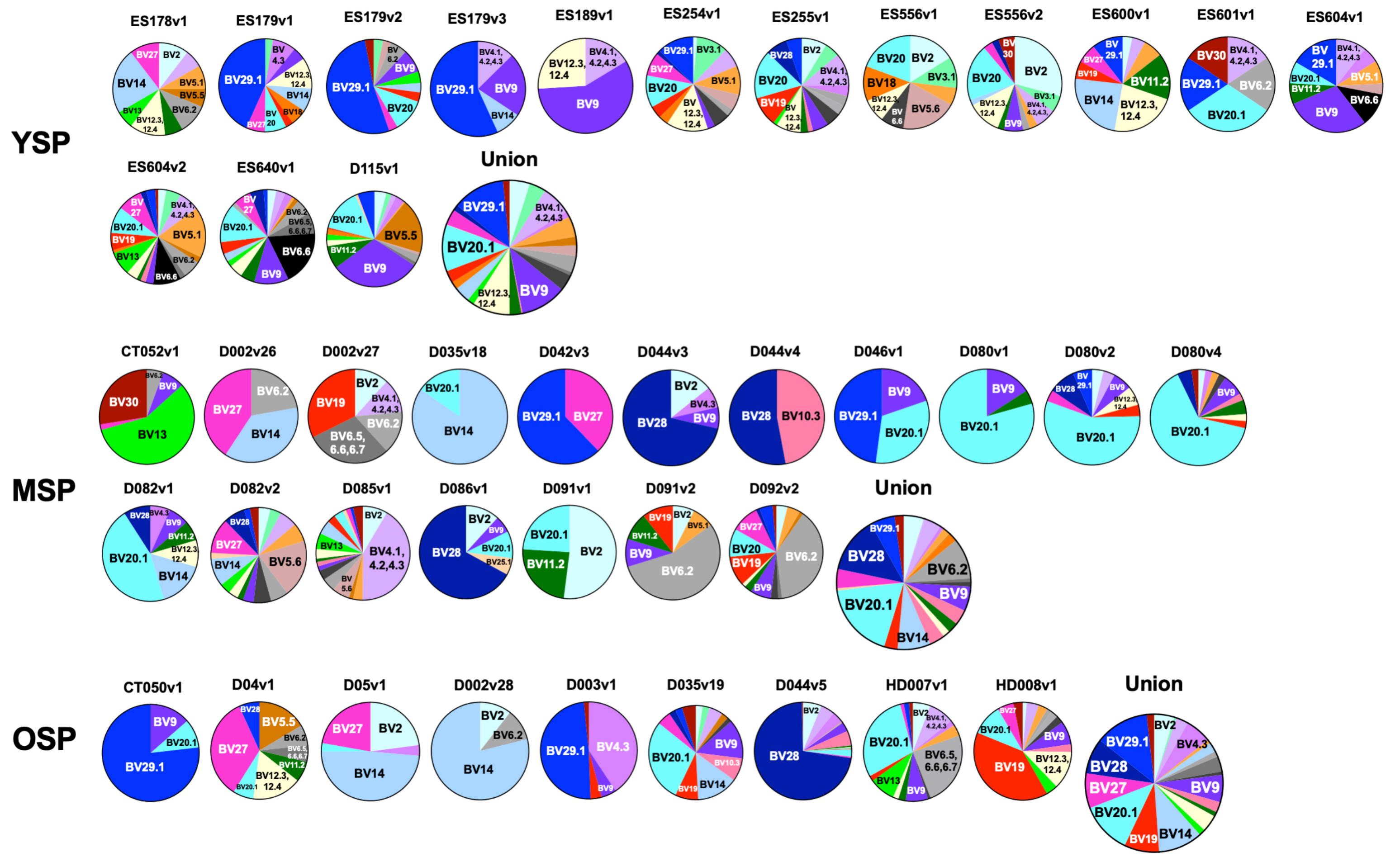

### Figure S4

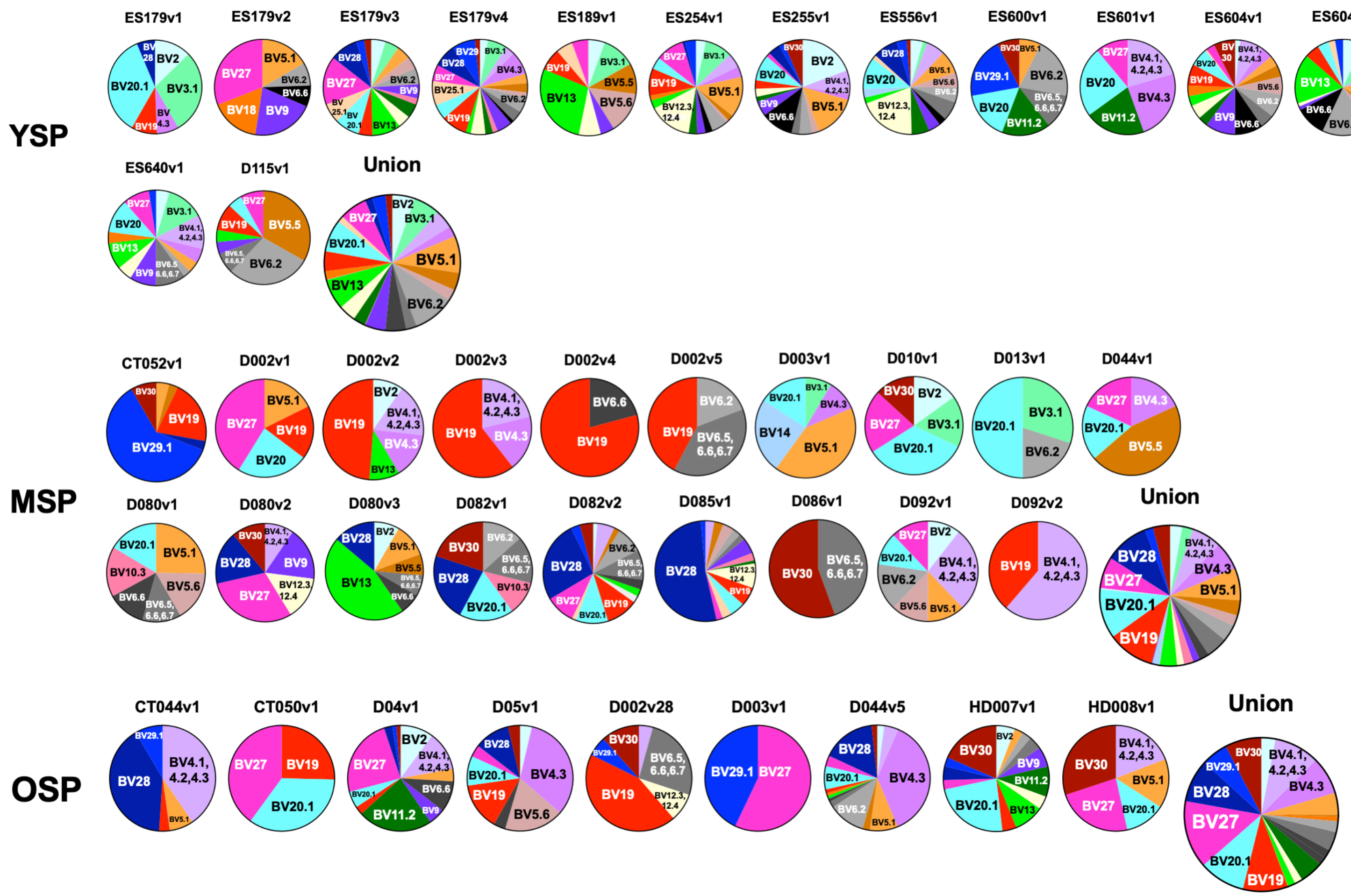

### Figure S5

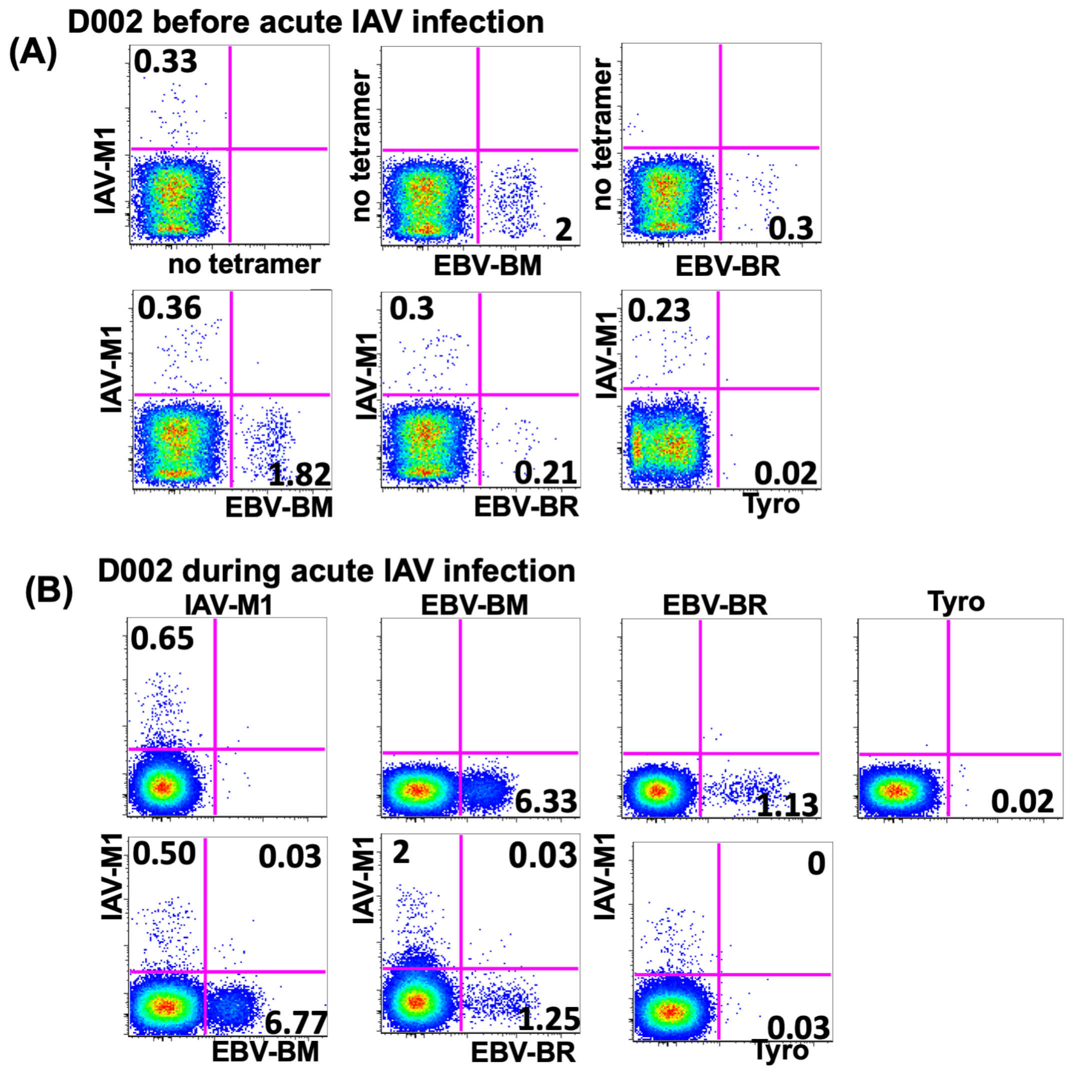
