## Supplementary material for "Co-evolution of human influenza A and Epstein Barr virus-specific CD8 ex vivo memory T cell receptor BV repertoires with increasing age": Table S1

### Supplemental Table 1: Characteristics of the study populations

| Study Population | Young EBV Sero-Positive (YSP) | Middle-age EBV Sero-positive (MSP) | Older EBV Sero- positive (OSP) |
|---|---|---|---|
| Median age in years (range) | 18.8 (18-22) | 37 (25-59) | 73.6 (60-93) |
| Number of subjects (n) | 25 | 13 | 11 |
| Sex (F/M) | 20/4 | 7/5 | 6/5 |
